# Evidence for Elton’s diversity-invasibility hypothesis from belowground

**DOI:** 10.1101/2020.03.11.987883

**Authors:** Zhijie Zhang, Yanjie Liu, Caroline Brunel, Mark van Kleunen

## Abstract

Sixty year ago, Elton proposed that diverse communities are more resistant to biological invasion. However, still little is known about which processes could drive this diversity-invasibility relationship. Here we examined whether plant-soil feedback on alien invaders is more negative when the soil originates from multiple native species. We trained soils with five individually grown native species, and used amplicon sequencing to analyze the resulting bacterial and fungal soil communities. We mixed the soils to create trained soils from one, two or four native species. We then grew four alien species separately on these differently trained soils. In the soil-conditioning phase, the five native species built species-specific bacterial and fungal communities in their rhizospheres. In the test phase, it did not matter whether the soil had been trained by one or two native species. However, the alien species achieved 11.7% less aboveground biomass when grown on soils trained by four native species than on soils trained by two native species. Our results showed for the first time, that plant-soil feedback could be a process that contributes to the negative relationship between diversity and invasibility.

## Introduction

In the last centuries, most regions of the world have been invaded by alien organisms (Dawson et al. 2017), and these invasions are still increasing (Pyšek et al. 2017, Seebens et al. 2018). The increasing numbers of naturalized alien species have stimulated discussion on how to increase community resistance to biological invasion. Elton (1958) proposed that diverse communities are more resistant to biological invasion. Support for Elton’s diversity-invasibility hypothesis has arisen from experiments, particularly on plants (Levine 2000, Levine et al. 2004). Most of them focused on the relationship between diversity-invasibility, but not on the underlying mechanism. Theoretical models usually ascribed this relationship to a lack of available resources in diverse communities (Case 1990, Byers and Noonburg 2003), likely because Elton (1958) introduced his hypothesis with several examples where resource competition was likely to determine invasibility. However, as already acknowledged by Elton (1958), resource competition is not the only determinant of invasibility. Other processes, such as plant-soil feedback, could also affect invasions (Klironomos 2002, Dawson et al. 2016), and could potentially drive the relationship between diversity and invasibility.

Plant-soil feedback refers to the process where plants influence the soil environment, particularly the soil biota, through inputs of organic matter and other chemical compounds, which influence performance of the plants that later grow on the same soil (Bever et al. 1997, Bennett and Klironomos 2019). In the last two decades, numerous studies have tested how single species affect growth of later plants through plant-soil feedback (Kulmatiski et al. 2008). However, only few studies tested how mixtures of multiple plant species affect later plants through plant-soil feedback (but see Müller *et al.*, 2015; Robin *et al.*, 2018 for how plant communities affects later plants). This is surprising, given that natural plant communities typically consist of multiple intermingled species. Consequently, we know little about whether diverse plant communities better resist later plants through plant-soil feedback than less-diverse communities do.

Several hypotheses offer insights into how plant-soil feedback could affect the diversity-invasibility relationship, but they predict different patterns. First, the amplification-effect hypothesis proposes that a diverse community of plant species harbors a greater diversity and abundance of pathogens (Hudson et al. 2006, Keesing et al. 2006). This increases the likelihood that some of those pathogens will negatively affect invaders. Following this logic, diverse communities should be better able to resist alien plants. Second, the dilution-effect hypothesis proposes that diverse communities reduce the abundance of high susceptible hosts (Schmidt and Ostfeld 2001, Ostfeld and Keesing 2012), and that this reduces the prevalence of pathogens. Following this logic, diverse communities should be less able to resist alien plants. Third, the enemy-release hypothesis proposes that alien plants are released from enemies, such as pathogens (Mitchell and Power 2003). Following this logic, even when the diversity of plant communities affects pathogens, it might not strongly affect resistance of the plant communities. Given the contrasting predictions, empirical tests are necessary to test whether and how plant-soil feedback contributes to the diversity-invasibility relationship.

Here, we conducted a plant-soil feedback experiment with five native plant species to condition the soils and four alien plant species to test the effects of the diversity of native species. All nine species are widespread in Germany and can be locally abundant. We first grew each of the five native species individually to train soils. We collected rhizosphere soil and used amplicon sequencing to analyze the bacterial and fungal soil communities. We then mixed soil samples from one, two or four native species, and grew one of the four alien species on the soil mixture. This procedure allowed us to test whether diversity of native species affects alien species through plant-soil feedback.

## Materials and Methods

### Study species

We conducted a greenhouse experiment in which we used five herbaceous species (*Dactylis glomerata, Leontodon autumnalis, Lotus corniculatus, Plantago media, Salvia pratensis*) that are native to Germany to condition the soil, and four alien herbs (*Epilobium ciliatum, Lolium multiflorum, Senecio inaequidens, Vicia villosa*) as test species. We used multiple aliens as test species to increase our ability to generalize the results (van Kleunen et al. 2014). The classification of the status of the nine species was based on the Floraweb database (Bundesamt für Naturschutz 2003). All species are widespread in Germany and can be locally abundant. So, the four alien species can be classified as naturalized (and probably invasive; *sensu* Richardson *et al.*, 2000) in Germany. The nine species mainly occur in grasslands and overlap in their distributions (Bundesamt für Naturschutz 2003), so that they are likely to co-occur in nature. Seeds of the native species were obtained from Rieger-Hofmann GmbH (Blaufelden-Raboldshausen, Germany), and seeds of alien species were obtained from the Botanical Garden of the University of Konstanz.

### Experimental setup

The experiment consisted of three steps. First, we had the soil-conditioning phase, in which we trained the soils by growing each of the five native species individually on the soils. Then we collected soil samples from the soil-conditioning phase, and created different soil-mixture treatments. These treatments were soil mixtures trained by one species, two species or four species. Finally, in the test phase, we grew one of the four alien species individually on the soil mixture and determined its biomass production. Details on each of these steps are given below.

### Soil-conditioning phase

On 18 or 27 June 2018, we sowed seeds of the five native species into trays (10cm × 10cm × 5cm) filled with potting soil (Topferde^@^, Einheitserde Co., Sinntal-Altengronau, Germany). The soil and seeds were not sterilized. Because we wanted the different species to be in similar developmental stages at the beginning of the experiment, we sowed the species at different times (Table S1), according to their germination speed known from previous experiments. We then placed the trays with seeds in a greenhouse under natural light condition, with a temperature between 18 and 25°C.

On 9 July 2018, we transplanted for each of the five species 140 seedlings individually into 1.5-L pots filled with 25% field soil, 37.5% nonsterilized sand and 37.5% nonsterilized vermiculate. The field soil, served as inoculum to provide a live soil biotic community, and had been collected from a grassland site in the Botanical Garden of the University of Konstanz (47.69°N, 9.18°E). The soil had been sieved through a 1cm mesh to remove plant material and large stones. We placed each of the 700 pots on its own plastic dish to preserve water and to avoid cross-contamination through soil solutions leaking from the pots. We replaced seedlings that died within two weeks after transplanting by new ones. The pots were randomly allocated to positions in four greenhouse compartments (23°C/18°C day/night temperature, no additional light), fertilized with an NPK water soluble fertilizer (Universol Blue^®^, Everris, Nordhorn, Germany) at a concentration of 1‰ m/v seven times (100ml fertilizer per pot per time), watered as needed, and randomized twice.

From 22 to 26 October 2018, we harvested soils from each pot by first cutting the shoot and then removing the roots from the soil by sieving it through a 5-mm mesh. We randomly selected 120 pots of soil for each of the five species (600 pots in total), and used them to create the different soil-mixture treatments.

### Soil-mixture treatments

To create a species-diversity gradient, we mixed soil samples from the soil-conditioning phase to create three treatment levels, which we call diversity-1, diversity-2 and diversity-4 (Fig. 1). In the diversity-1 treatment, we collected 160 ml of soil from four different pots of the same species (40 ml from each pot). In the diversity-2 treatment, we collected 160 ml of soil from four different pots of two species (two pots per species). In the diversity-4 treatment, we collected 160 ml of soil from four different pots of four species (one pot per species). Then, the 160 ml of soil was mixed with 220 ml nonsterilized sand and 220 ml nonsterilized vermiculite, and put into 0.6-L pots, which were then used in the test phase described below. As we had a total of five native species in the soil-conditioning phase, this resulted in five possible species or combinations for diversity-1 and for diversity 4, but ten possible combinations for diversity-2. To have equal numbers of combinations for each diversity treatment, we therefore used only five out of the ten possible combinations for diversity-2. We did this in such a way that each species was included in two of the five combinations. Consequently, we had five species or species combinations for each diversity treatment. Each of those combinations was replicated eight times, such that we had a total of 120 pots for the test phase. Once we had collected soil from a pot of the soil conditioning phase, we did not use that pot again. This was done to avoid pseudoreplication (Hurlbert 1984), because this way no two pots in the test phase shared soil from the same pot of the soil-conditioning phase (i.e. they were independent).

**Figure 1.**
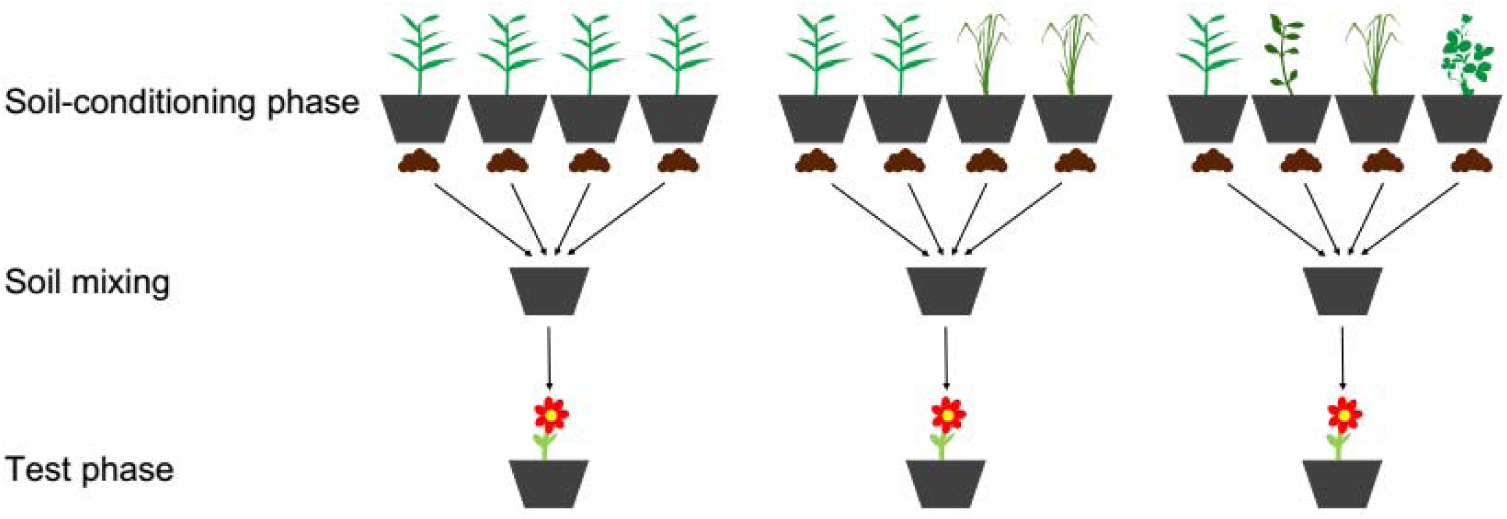
Graphical illustration of the experimental design. Soil samples trained by one, two or four native species were collected from four individuals (soil-conditioning phase). Then, the soil samples were used as inoculum, and mixed with sand and vermiculite (soil mixing). In each pot, one plant of each alien species was grown (test phase). The total amount of soil used to inoculate each pot was constant. Five native species and four alien species were used in the soil-conditioning phase and test phase, respectively.

In the diversity-1 treatment, we used soil of one single plant species, and in the diversity-2 treatment, we used soil of two plant species. Nevertheless, we still collected the soil from four different pots, just like in the diversity-4 treatment. We did this because recent studies argued that the approach of mixing soil samples *per se* can affect plants by increasing diversity of soil biota (Reinhart and Rinella 2016, Rinella and Reinhart 2018). More specifically, the composition of the soil biotic community can vary substantially across centimeters given their immense diversity (Decaëns 2010). So, microbes from different soil samples could differ in their identity, even when trained by the same plant species. This means that mixing soil samples might increase the diversity of mutualists and/or pathogens. Therefore, by consistently mixing soil samples from four pots for all diversity treatments, we reduced the potential side effects from soil mixing. In addition, we tested in a side experiment whether the mixing of soil samples from different individuals of the same species affects the strength of plant-soil feedback. We found no significant effect (Supplement S1), which indicates that the approach of mixing soil samples *per se* did not affect the relationship between diversity and invasibility.

### Test phase

Between 9 and 18 October 2018 (Table S1), we sowed the four alien species into trays filled with the same type of potting soil as in the soil conditioning phase. The soil and seeds were not sterilized. On 27 October 2018, we transplanted the seedlings into the 0.6-L pots prepared in the soil mixture phase. We assigned the seedlings in such a way that each alien species was replicated twice on each soil of a native species or native species combination (totaling 50 pots per test species). We placed each of the 200 pots on its own plastic dish, and they were randomly allocated to position in a greenhouse compartment (20°C/15°C day/night temperature, 14 h/10 h day/night light). The plants were fertilized with a water soluble NPK fertilizer (Universol Blue^®^, Everris, Nordhorn, Germany) at a concentration of 1‰ m/v four times (100ml fertilizer per pot per time), and their positions were re-randomized once. By homogenizing the soils in the soil-mixture treatments, inoculating a small amount (26.7% v/v) of trained soil and fertilizing the plants throughout the experiment, we were likely to remove plant-soil feedback differences due to abiotic properties. Therefore, any differences, if found, would be mainly due to differences in biotic properties of the soils. On 18 December 2018, seven weeks after the start of the test phase, we harvested all aboveground plant parts, and then washed the roots free from soil. The biomass was dried at 70°C to constant weight, and weighed to the nearest milligram.

### Soil sampling, DNA extraction, amplicon sequencing and bioinformatics

From 22 to 26 October 2018, when we harvested the trained soil, we randomly selected six pots for each of the five native species. For each of these pots, 10-20ml rhizosphere-soil was collected into sterile 50-ml cylindrical tube, and stored at −80□ until required for DNA extraction. We extracted DNA from 0.25g of each soil sample using the DNeasy® PowerSoil® Kit (Qiagen, Hilden, Germany), following the manufacturer’s protocol.

PCR amplifications and amplicon sequencing were then performed by Novogene (Beijing China). In brief, V3-V4 region of bacterial 16S rDNA gene was amplified in triplicate with the universal primers 341F/806R (forward primer: 5’- CCTAYGGGRBGCASCAG-3’; reverse primer: 5’- GGACTACNNGGGTATCTAAT- 3’)(Klindworth et al. 2012). The ITS2 region of fungal rDNA gene was amplified in triplicate with the primers specific to this locus (forward primer: 5’- GCATCGATGAAGAACGCAGC-3’; reverse primer: 5’- TCCTCCGCTTATTGATATGC- 3’)(Orgiazzi et al. 2012). All PCR reactions were carried out with Phusion® High-Fidelity PCR Master Mix (New England Biolabs). PCR products were mixed in equidensity ratios. Then, mixtures of PCR products were purified with Gel Extraction Kit (Qiagen, Germany). The libraries were generated with NEBNext® Ultra™ DNA Library Prep Kit for Illumina and analyzed using the Illumina platform.

We processed the raw sequences with the *DADA2* pipeline (Callahan et al. 2016), which is designed to resolve exact biological sequences (Amplicon Sequence Variants [ASVs] or phylotypes) from Illumina sequence data and does not involve sequence clustering (Callahan et al. 2017). The detailed process was described in Brunel et al. (2019). In short, we removed primers and adapter with the *cutadapt* package (Martin 2011), merged paired-end sequences, and removed chimeras. Then, we determined taxonomy assignments against derivative reformatting of the UNITE (Nilsson et al. 2018) and the SILVA (Quast et al. 2013) taxonomic databases for fungi and bacteria, respectively. Last, we rarefied bacteria and fungi to 30,000 and 10,000 reads, respectively, to account for differences in sequencing depths. One sample that did not have enough reads for bacteria and two samples that did not have enough PCR product for fungi were excluded from soil analyses. Putative fungal functional groups (e.g. Arbuscular mycorrhiza fungi, plant pathogens and endophytes) were identified using FUNGuild (Nguyen et al. 2016).

### Statistical analyses

To test whether plant-soil feedback on alien invaders is more negative when the soil originates from multiple native species, we used mixed effect models to analyze the biomass production of the alien plants, as implemented in the *nlme* package (Pinheiro et al. 2018) with R 3.4.0 (R Core Team 2017). This analysis was restricted to the subset of 120 plants that grew in pots inoculated with soil samples from four different pots of the conditioning phase (Fig. 1). The models included aboveground, belowground or total biomass as the response variables, and soil-diversity treatment (i.e. diversity-1, −2 and −4) as the fixed effect. The models included identity of the alien species and identity of the soil (i.e. identity of the native species for diversity-1, and identity of the species combination for diversity-2 and −4 treatments) as random effects. Because initial data exploration showed that the effect of diversity is nonlinear, we included soil-diversity as a categorical instead of as a continuous variable. To improve homoscedasticity of residuals, we allowed the alien species and soil-diversity treatments to have different variances by using the *varComb* and *varIdent* function (Zuur et al. 2009). We performed multiple pairwise comparisons with Tukey correction to test for differences among the three soil-diversity levels with the *multicomp* package (Hothorn et al. 2008).

To test whether the five native plant species differently trained the soil microbial communities, we analyzed whether alpha and beta diversities were related to plant species identity. As alpha diversities, we calculated both species richness and Shannon indexes, which were analyzed with linear models. The models included plant species identity as the explanatory variables. As the beta diversity, we calculated Bray–Curtis dissimilarities using reads relative abundances, which were analyzed with permutational analysis of variance (PERMANOVA), as implemented in the *adonis* function of the *vegan* package (Oksanen et al. 2019), again including plant species as explanatory variable. We used nonmetric multidimensional scaling (NMDS) to illustrate differences in the soil microbioal communities of the plant species. We determined shared and unshared taxa among plant species (based on phylotypes occurring in at least one of the samples) and visualized them using 5 set-Venn diagrams with the *eulerr* package (Larsson 2018).

## Results

### Microbial communities of trained soils

We detected a total of 21,724 bacterial phylotypes and 1,231 fungal phylotypes. Identity of native plant species did not significantly explain Shannon diversity of (Table S4; Fig. S4) and variation in the composition of fungal communities (PERMANOVA, *r*^*2*^ = 0.179, *F* = 1.251, *p* = 0.144, Fig. 2b). However, identity of native plant species marginally explained species richness of bacterial and fungal communities (Table S4; Fig. S4), and largely explained the variation in composition of bacterial communities (PERMANOVA, *r*^*2*^ = 0.274, *F* = 2.266, *p* < 0.001, Fig. 2a). In addition, 55.1% of bacterial phylotypes and 61.4% of fungal phylotypes were not shared among native plant species (Fig. 3c&d). Fungal functional groups (i.e. AMF, plant pathogen and endophyte) could be assigned to 495 phylotypes, representing 34.0 % of ITS sequence reads. Of these phylotypes, 23 were identified as AMF, 103 as endophytes and 136 as plant pathogens (54 phylotypes were identified as both plant pathogen and endophyte). Variation in composition of identified endophytes and plant fungal pathogens was not significantly explained by identity of the native plant species (endophyte: PERMANOVA, *r*^*2*^ = 0.172, *F* = 1.194, *p* = 0.234; pathogen: PERMANOVA, *r*^*2*^ = 0.166, *F* = 1.143, *p* = 0.177; Fig. S5; for AMF we had insufficient data).

**Figure 2.**
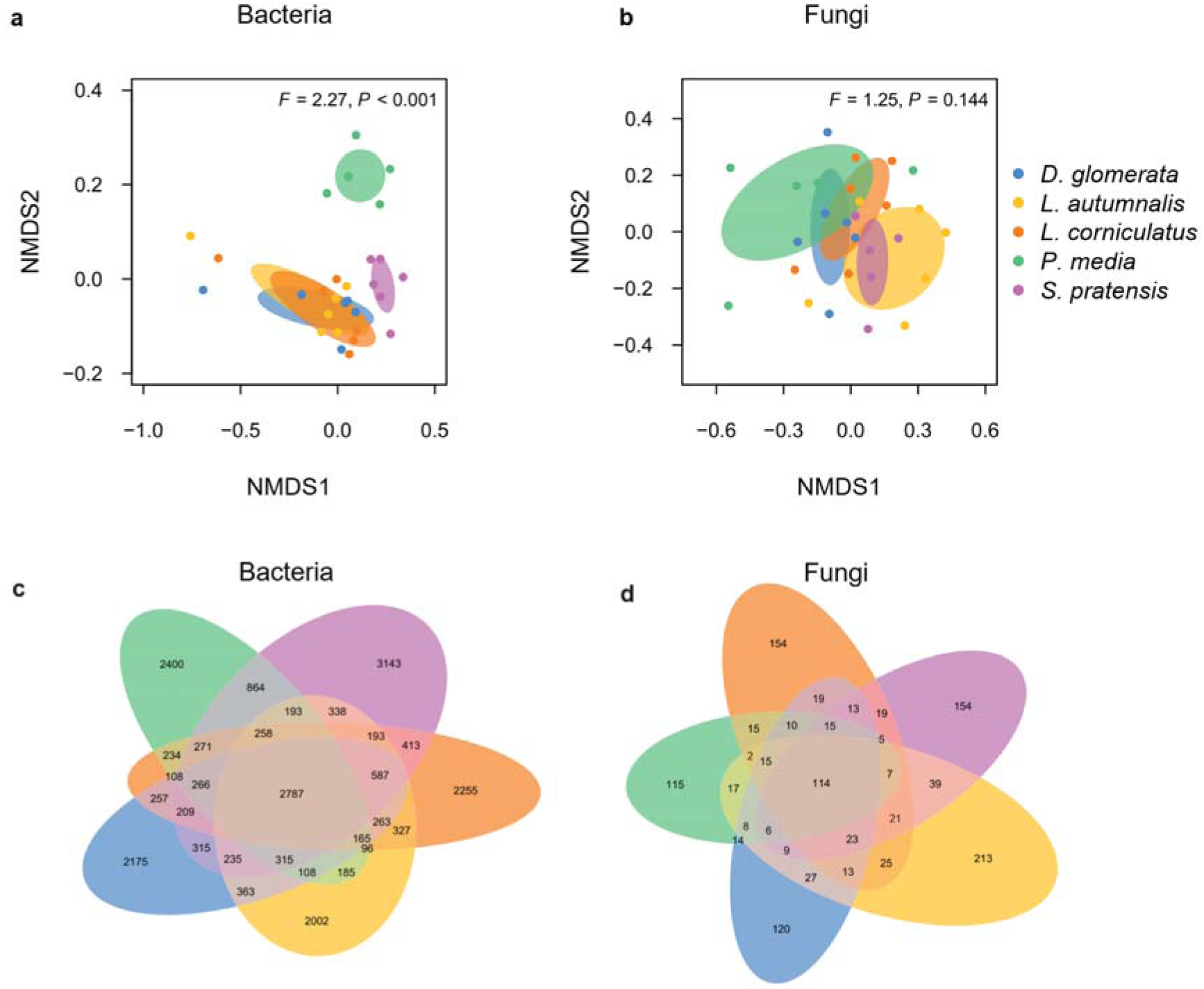
Dissimilarity (beta diversity and unshared phylotypes) of bacterial (a, c) and fungal (b, d) community composition among soils trained by different native species. Different colors represent different species. In the NMDS figures (upper panel), data points represent soil samples. Ellipses represent means ± 1 SDs for each native plant species.

**Figure 3.**
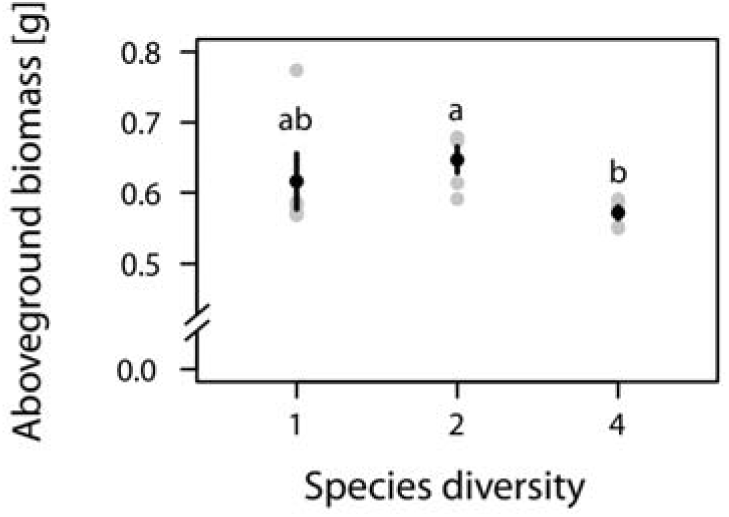
Aboveground biomass (means ± SEs) of alien plants when their pots had been inoculated with soil trained by one, two or four native species. Grey dots represent mean values of aboveground biomass of alien plants when grown on soil trained by different native species or by different native species combinations. Different letters above the error bars indicate significant differences (P < 0.05) between different treatments based on Tukey’s multiple comparison.

### Biomass of plants in the test phase

Diversity of the native species used to create the soil inocula affected the aboveground biomass production of the alien plants significantly (χ^2^ = 7.956, *P* = 0.019; Table S2). The aboveground biomass of the alien plants was not significantly different between the diversity-1 and diversity-2 treatments (Fig. 3; diversity-2 *vs* diversity-1, z = 1.123, *P* = 0.500). In other words, it did not matter whether the soil inoculum came from one or two native species. However, the alien plants produced 11.7% less aboveground biomass in the diversity-4 treatment than in the diversity-2 treatment (Fig. 3; diversity-4 *vs* diversity-2, z = −2.964, *P* = 0.008). Although the alien plants tended to achieve also less aboveground biomass (−7.26%) in the diversity-4 treatment than in the diversity-1 treatment (Fig. 3), the difference was not statistically significant (diversity-4 *vs* diversity-1; z = −1.638, *P* = 0.230). A similar pattern was found for total biomass, but there the effect of diversity-4 *vs* diversity-2 was marginally significant (Fig. S3a; diversity-4 *vs* diversity-2, z = −2.067, *P* = 0.096). Belowground biomass did not differ among the three diversity treatments (Fig. S3b; Table S2).

## Discussion

We here show that alien plants achieved less aboveground biomass when grown on a mixture of soil trained by four native species than when grown on a mixture of soil trained by two native species. This suggests that diverse native communities could impede plant invasion through plant-soil feedback. So, whereas previous studies frequently ascribed the negative relationship between diversity and invasibility to resource competition (Byers and Noonburg 2003), we showed for the first time that this relationship could also be driven by plant-soil feedback.

The negative effect of soil from diverse native communities on alien plants could be mediated by the establishment of a diverse community of soil pathogens during the conditioning phase. This is because we found that different native species built species-specific bacterial and fungal communities in their rhizospheres. Consequently, diverse plant communities harbored a greater diversity of soil microbiota, many of which might be plant pathogens with a particularly strong negative impact on the alien plant. This finding thus supports the amplification-effect hypothesis (Keesing et al. 2006), and does not support the predictions of the dilution-effect hypothesis (Schmidt and Ostfeld 2001, Ostfeld and Keesing 2012) and the enemy-release hypothesis (Mitchell and Power 2003). One explanation for the latter could be that, although alien plants could escape from their co-evolved enemies (i.e. specialist enemies), they might still encounter biotic resistance from generalist enemies (Maron and Vilà 2001, Dawson et al. 2014, Zhang et al. 2018). This explanation becomes more plausible given that the dilution effect is less likely to happen when pathogens are generalists (Power and Mitchell 2004). Therefore, our finding that diverse communities are more resistant to biological invasion might be driven by generalist pathogens.

Although diverse native communities could suppress alien plant performance through plant-soil feedback, this effect was only significant when we compared the alien plants grown on soils trained by four native species with those grown on soils trained by two native species. The comparisons with plants grown on soils trained by only one native species (i.e. diversity-1 *vs* diversity-2, and diversity-1 *vs* diversity-4) were not significant. Probably, this is because at low diversity, the variation among native species in their effects on the alien species was large (see error bars of diversity-1 in Fig. 3), which limited the statistical power. Nevertheless, the alien plants on soil trained by four native species tended to produce less aboveground biomass than the alien plants grown on soil trained by one native species. Moreover, invasion is a probabilistic process (Crawley 1987, Levine and D’Antonio 1999), and the probability that a community includes native species that can impede alien plants increases with the diversity of the community. Such a so-called selection effect (Huston 1997) is also indicated by the 14.9% higher variance in the diversity-1 treatment than diversity-2 and −4 treatments. If most native species do not strongly affect the alien plants, it will, however, be difficult to detect a diversity effect at low diversities. This could explain the absence of a difference in biomass production of aliens between the diversity-1 and diversity-2 treatments.

Unexpectedly, root biomass of alien plants did not change with diversity of native communities. As roots are in direct contact with soil, effects of plant-soil feedback are expected to first emerge in the roots and then subsequently affect the shoots. Possibly, roots in all diversity treatments were limited by pot size. Total plant biomass per unit pot volume was 1.45 g/L on average, which is lower than most pot experiments but higher than the threshold (1 g/L) where pot limitation starts to happen (Poorter et al. 2012). Alternatively, our finding may reflect that the alien plants have high phenotypic plasticity (Davidson et al. 2011), and changed their root-shoot allocation in such a way that they could maintain maximum root biomass under different conditions.

### Future directions

Invasibility is defined as invasion growth rates of invaders, that is, population growth rates when the invader is introduced into native plant communities at low densities (Case 1990). Therefore, it will be more realistic and straightforward to include competition from native plants in the test phase. However, invasion growth rate can be decomposed into two parts (see Cardinaux, Hart & Alexander, 2018 for a mathematical expression): 1) intrinsic growth rate of the invader, i.e. growth rates when grown without competition; 2) competitive effect from the native community. As intrinsic growth rates of aliens decreased when soil were conditioned by four native species in our study, invasion growth rates of the aliens would decrease accordingly. Still, it remains an open question whether the competitive effect from the native community depends on the diversity of native species used to condition the soil. Besides, while Elton (1958) introduced the diversity-invasibility hypothesis with several examples on resource competition, he acknowledged that other processes, such as enemy-mediated apparent competition, can also determine invasibility. Because different processes can be correlated with diversity of native plants, testing multiple processes together will offer more insights.

We found that the diversity-4 treatment reduced biomass of alien species by 11.7%. It remains unknown whether this reduction is sufficient to eliminate the dominance of alien species over natives. Previous multispecies experiments offer us some clues. They showed that native species produced 13.6∼38.3% less biomass than aliens (Godoy et al. 2011, Zhang and van Kleunen 2019). Therefore, the dominance of alien species over natives would be alleviated if native species are less limited by the negative soil feedback from diverse native communities than aliens are. As we know little about whether native and alien species respond differently to soil feedback from diverse native communities, direct tests that include both alien and native species in the test phase are needed.

We measured the plants in their juvenile life stage. However, establishment of aliens into native communities also includes earlier stages, such as seed germination, and later stages, such as reproduction. As a recent experiment showed that the magnitude and direction of plant-soil feedback can vary across different life stages (Dudenhöffer et al. 2018), more comprehensive tests require also longer-term investigations that consider all life stages.

The soil that we used to create the different diversity levels came from pots in which plants were grown without competition. It could be argued that the lack of direct plant-plant interactions in the soil-conditioning phase could have biased our results. For example, competition can affect plant growth and possibly root exudation, which may change the microbial community, and thus the magnitude or even direction of plant-soil feedback in the test phase. By mixing soil trained by different species rather than directly using soil trained by communities that consist of multiple species, we removed the effect of plant competition on the microbial community. Nevertheless, our method allowed us to isolate the effect of diversity that resulted from plant-soil feedback. Still, more research is needed to identify whether and how plant competition modifies diversity effect of plant-soil feedback.

## Conclusions

Sixty years after Elton (1958) proposed the diversity-invasibility hypothesis, mounting evidence has shown that, at the local scale, diverse communities are better at resisting invasion by alien plants. There was already evidence showing that competition is likely to contribute to this relationship (Byun et al. 2013, Feng et al. 2018). However, here we show for the first time, that plant-soil feedback might also contribute to the negative diversity-invasibility relationship. An important next step would be to test the relative importance of plant-soil feedback and other processes, such as plant competition, in determining the diversity-invasibility relationship.

## Supporting information

all supplements

## Acknowledgements

We thank O. Ficht, M. Fuchs, S. Gommel, E. Mamonova, V. Pasqualetto, C. Rabung, B. Rüter, H. Vahlenkamp and E. Werner for practical assistance; and X. Liu for comments on a previous version. ZZ acknowledges funding from the China Scholarship Council (201606100049) and support from the International Max Planck Research School for Organismal Biology. The authors declare no conflict of interest. YL acknowledges funding from the Chinese Academy of Sciences (Y9H1011001).

## Author contributions

ZZ, YL and MvK designed the experiment. ZZ and YL performed the experiment. CB led the soil analyses. ZZ analyzed the data and led the writing with input from all others.

## Data accessibility

Should the manuscript be accepted, the data supporting the results will be archived in an appropriate public repository (Dryad, Figshare or Hal) and the data DOI will be included at the end of the article.

