## Supplementary material for "Evidence for Elton’s diversity-invasibility hypothesis from belowground": all supplements

### **Supplement S1** A side experiment testing whether the mixing of soil samples from different individuals of same species affects the strength of plant-soil feedback.

Introduction

Although the potential side effects of mixing soil samples have been recognized by many studies (e.g. Cahill Jr *et al.*, 2017), few studies empirically tested them (Gundale et al. 2019). To explicitly test whether the mixing of soil samples from different individuals of the same species affects the strength of plant-soil feedback (i.e. affects growth of plants in the test phase), we added two further one-species (i.e. diversity-1) treatments (Fig. S1). In one of the treatments, we collected 160 ml of soil from just one pot, and in the other treatment, we collected a total of 160 ml soil from two different pots (80 ml from each pot). Each of these two additional treatments was replicated eight times for each of the five native species, such that we had a total of 80 additional pots for the test phase.

Method

To test whether mixing soil samples affected growth of alien species, we used a similar analysis approach as for the test of soil-diversity effects. The only differences were that we used the subset of 120 alien plants that grew on soils from single species (i.e. diversity-1; Fig. S1), and that we used the number of soil samples instead of soil-diversity as the fixed effect.

Results

We found that the approach of mixing soil samples *per se* did not affect the biomass production of the alien plants (Table S3). When the alien plants grew on soils from a single native species, their aboveground biomass did not change with the number of pots from which the soil had been collected (Fig. S2; Table S3; 4 vs 2 soil samples, z = -0.296, *P* =0.953; 4 vs 1 soil samples, z = 0.228, *P* =0.972; 2 vs 1 soil samples, z = 0.519, *P* =0.862). Similar patterns were found for total and belowground biomass (Fig. S3c & S3d; Table S3).

Discussion

When growing alien plants on soil from a single native species, we found that aboveground biomass of the alien plants did not change with the number of individuals (i.e. pots) from which the trained soil came. It is worth noting, however, that, in our study, genetic variation among the individuals of a native species was likely to be low, because the seeds used for each species came from the same location. In addition, the field where we collected the soil to provide live soil biota is relatively homogeneous. Possibly, if genetic variation of the species used for soil conditioning would be high or if soil biota varies substantially across small spatial scales, it might matter whether one mixes soil samples from different individuals or not. Nevertheless, in our study, it is unlikely that the approach of mixing soil samples *per se* affected the relationship between diversity and invasibility.

### **Table S1** The five native and four naturalized alien perennial herbs included in the study.

| **Species** | **Family** | **status** | **Sowing date** |
| --- | --- | --- | --- |
| **Soil-conditioning phase** |  |  |  |
| *Dactylis glomerata* | Poaceae | native | Jun-27 2018 |
| *Leontodon autumnalis* | Asteraceae | native | Jun-25 2018 |
| *Lotus corniculatus* | Fabaceae | native | Jun-18 2018 |
| *Plantago media* | Plantaginaceae | native | Jun-25 2018 |
| *Salvia pratensis* | Lamiaceae | native | Jun-25 2018 |
| **Test phase** |  |  |  |
| *Epilobium ciliatum* | Onagraceae | alien | Oct-9 2018 |
| *Lolium multiflorum* | Poaceae | alien | Oct-18 2018 |
| *Senecio inaequidens* | Asteraceae | alien | Oct-16 2018 |
| *Vicia villosa* | Fabaceae | alien | Oct-16 2018 |

### **Table S2** Effects of native diversity on biomass production of alien plant species.

Significant (*P* < 0.05) effects are shown in bold. The significance of fixed effects in the mixed models was assessed with likelihood-ratio tests when comparing models with and without the effect of interest.

|  | Aboveground | | |  | Belowground | |  | Total | |
| --- | --- | --- | --- | --- | --- | --- | --- | --- | --- |
|  | χ^2^ | *P* | |  | χ^2^ | *P* |  | χ^2^ | *P* |
| Native diversity | 7.956 | **0.019** | |  | 0.125 | 0.939 |  | 4.295 | 0.117 |
| *Random effects* | SD | |  |  | SD |  |  | SD |  |
| Identity of alien | 0.064 | |  |  | 0.140 |  |  | 0.206 |  |
| Identity of soil | 0.000 | |  |  | 0.000 |  |  | 0.000 |  |
| Residual | 0.185 | |  |  | 0.061 |  |  | 0.233 |  |

### **Table S3** Effects of soil-samples number on biomass production of alien plant species. The significance of fixed effects in the mixed models was assessed with likelihood-ratio tests when comparing models with and without the effect of interest.

|  | Aboveground | | |  | Belowground | |  | Total | |
| --- | --- | --- | --- | --- | --- | --- | --- | --- | --- |
|  | χ^2^ | *P* | |  | χ^2^ | *P* |  | χ^2^ | *P* |
| Number of soil samples | 0.345 | 0.842 | |  | 0.725 | 0.696 |  | 0.161 | 0.923 |
| *Random effects* | SD | |  |  | SD |  |  | SD |  |
| Identity of alien | 0.063 | |  |  | 0.122 |  |  | 0.176 |  |
| Identity of soil | 0.062 | |  |  | 0.030 |  |  | 0.103 |  |
| Residual | 0.146 | |  |  | 0.064 |  |  | 0.184 |  |

### **Table S4** Effects of native species identity on alpha diversity (species richness and Shannon diversity) of bacterial and fungal communities.

Bateria:

|  | Species richness | | | | | Shannon diversity | | |
| --- | --- | --- | --- | --- | --- | --- | --- | --- |
|  | Sum of sq | RSS | *P* |  | Sum of sq | | RSS | *P* |
| Species | 3030212 | 12629711 | 0.093 |  | 0.048 | | 0.387 | 0.431 |

Fungi:

|  | Species richness | | | | | Shannon diversity | | |
| --- | --- | --- | --- | --- | --- | --- | --- | --- |
|  | Sum of sq | RSS | *P* |  | Sum of sq | | RSS | *P* |
| Species | 5701 | 22926 | 0.091 |  | 1.304 | | 7.121 | 0.226 |

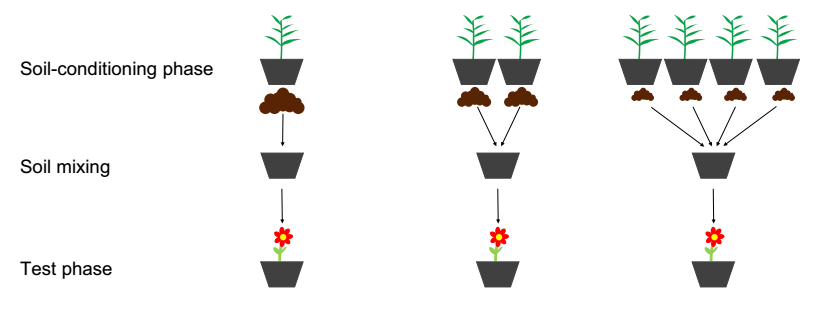

### **Figure S1** Graphical illustration of the experimental design for testing the effect of mixing soil, soil samples trained by one of the five native species were collected from one, two or four individuals (soil-conditioning phase). Then, the soil samples were used as inoculum, and mixed with sand and vermiculate (soil-mixing). In each pot, one plant of each alien species was grown (test phase). The total amount of soil used to inoculate each pot was constant. Five native species and four alien species were used in the soil-conditioning phase and test phase, respectively.

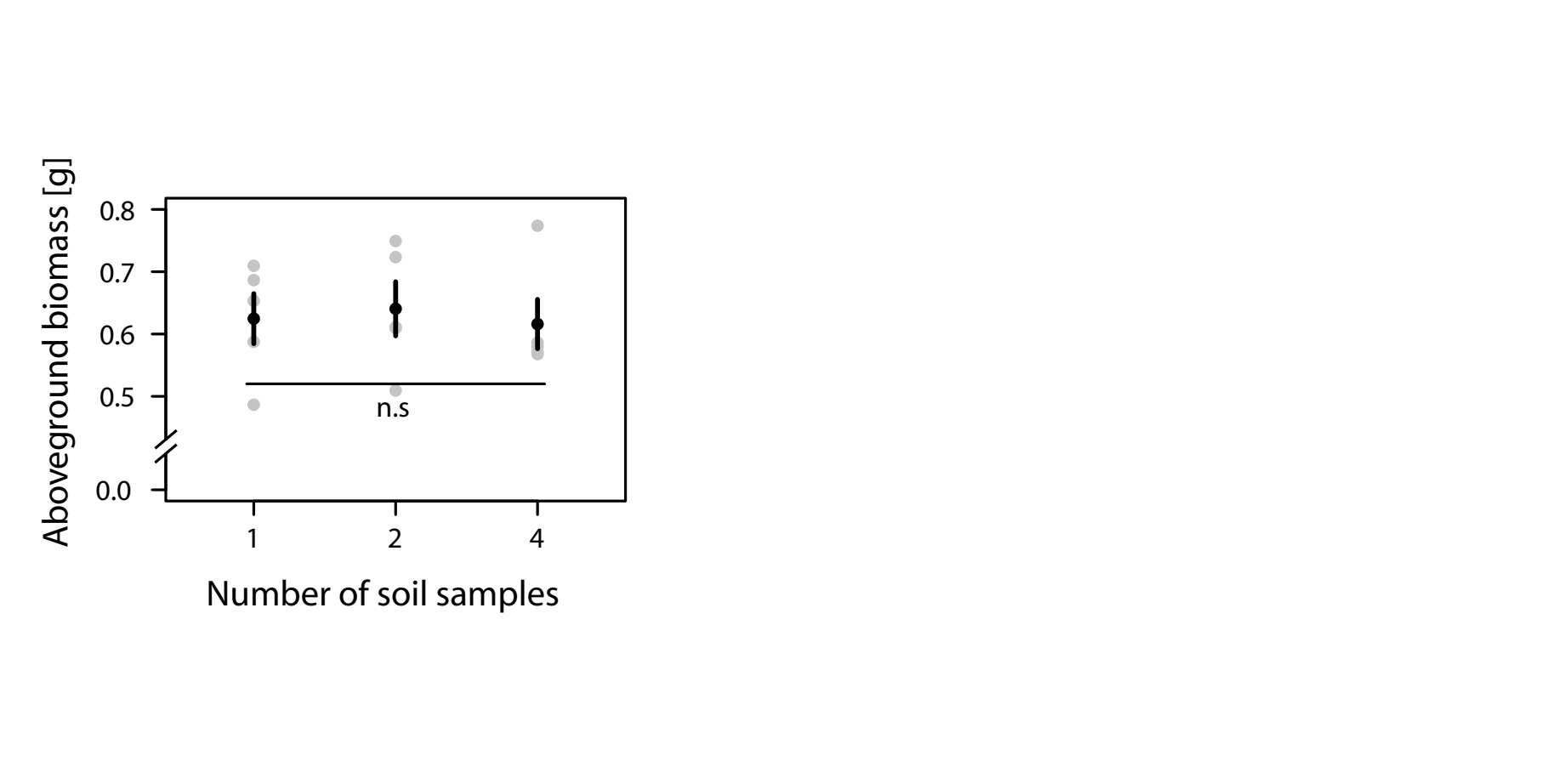

### **Figure S2** Aboveground biomass (means ± SEs) of alien plants when their pots had been inoculated with soil samples trained by one, two or four individuals of the same species. Grey dots represent mean values of aboveground biomass of alien plants when grown on soil trained by different native species. n.s. indicates no significant differences among treatments.

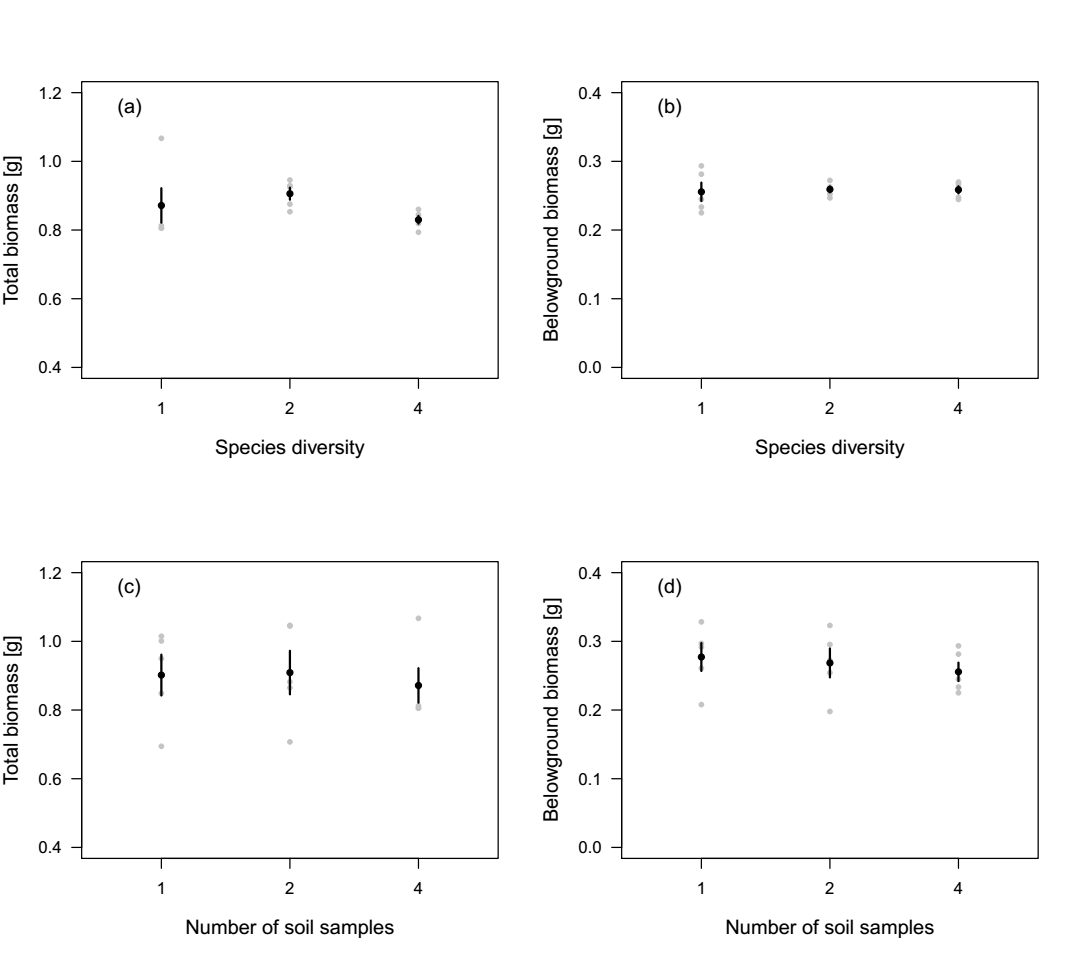

### **Figure S3** Total biomass (a, c) and belowground biomass (b, d) (means ± SEs) of alien plants (a, b) when their pots were inoculated with soil trained by one, two or four native species, and (c, d) when their pots were inoculated with soil trained by one, two or four individuals of the same species. Grey dots represent mean values of belowground or total biomass of alien plants when grown on soil trained by different native species or by different native species combinations. Tukey’s multiple comparisons showed no significant differences among treatments.

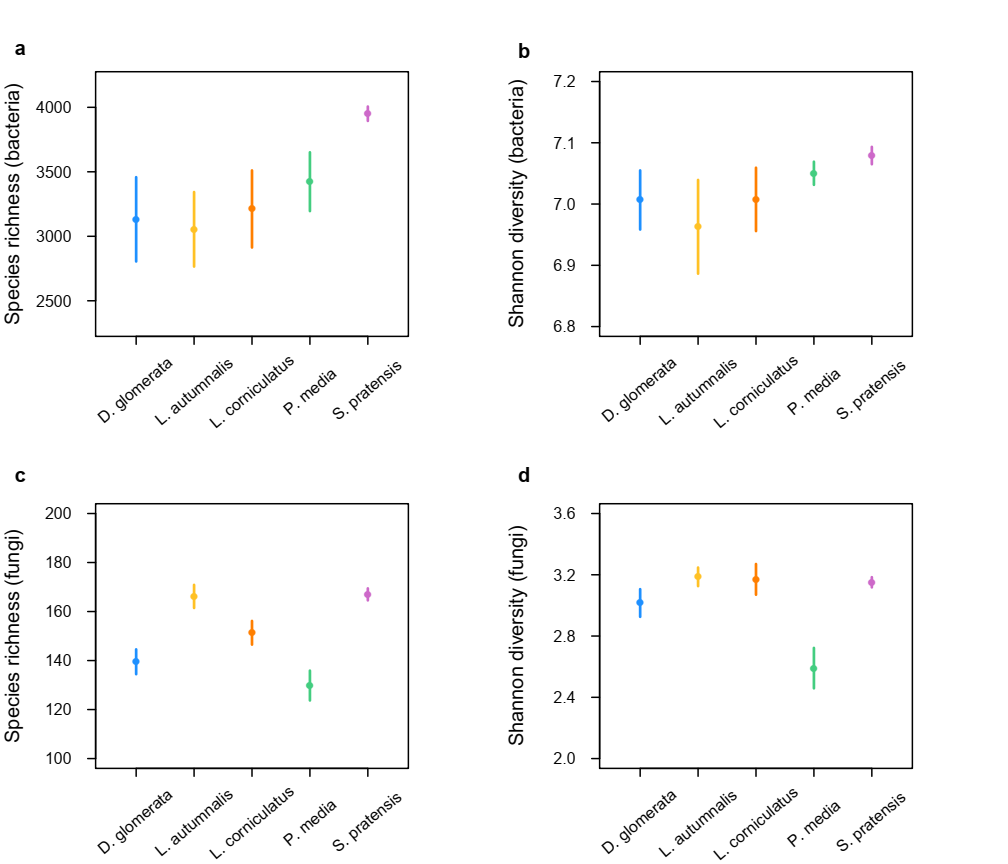

### **Figure S4** Species richness and Shannon diversity of bacterial (a, b) and fungal (c, d) communities in soils trained by different native species. Means ± SEs are presented.

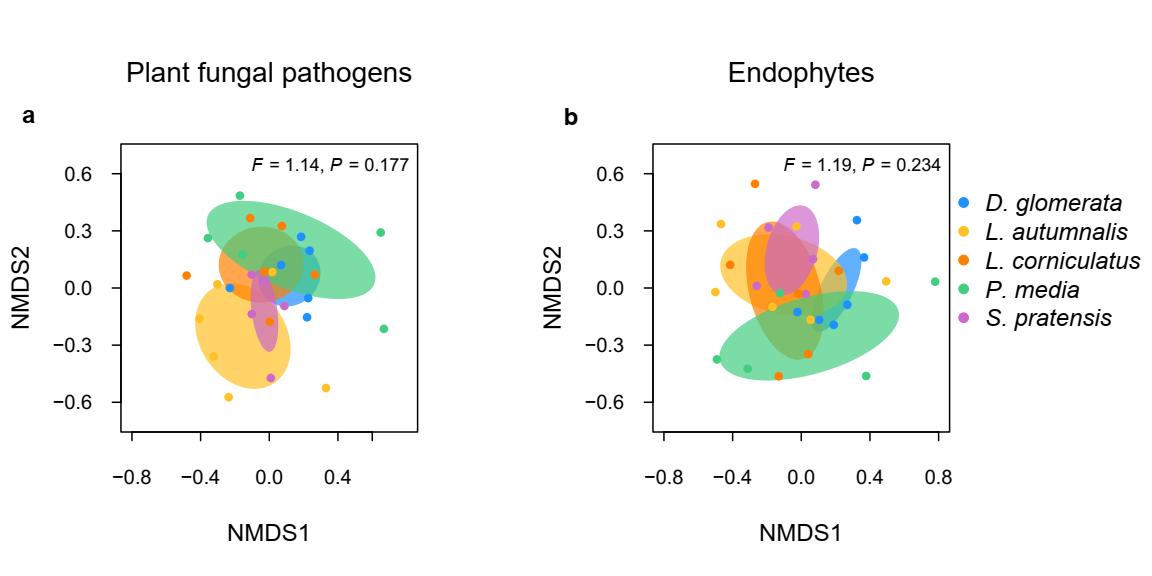

### **Figure S5** Dissimilarity (beta diversity) of plant fungal pathogens (a) and endophytes (b) community composition among soils trained by different native species. Data points represent soil samples. Ellipses represent means ± 1 SDs for each native plant species. Dissimilarity of AMF is not shown due to insufficient data.

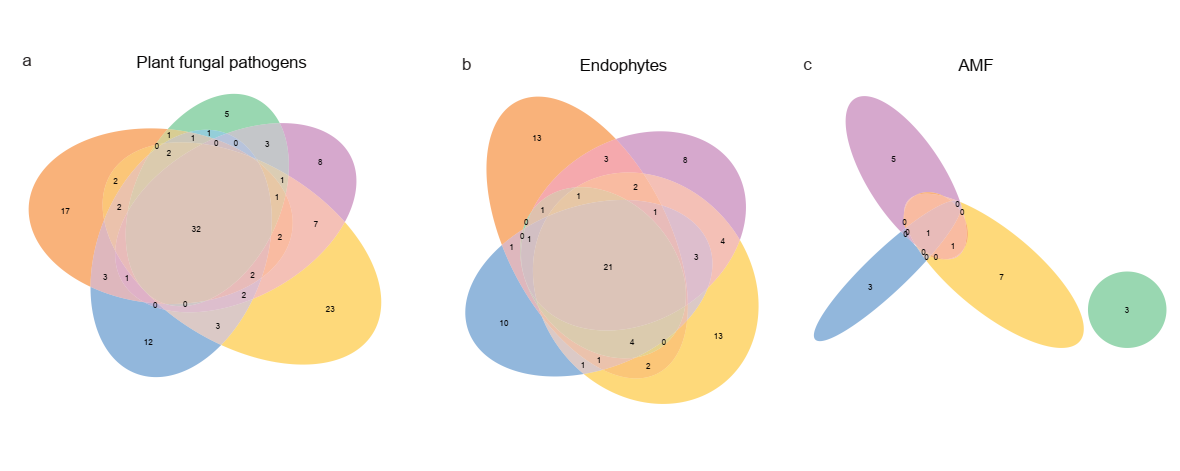

### **Figure S6** Dissimilarity (unshared phylotypes) of plant fungal pathogens (a) and endophytes (b) and AMF (c) community composition among soils trained by different native species.
